# Myotube hypertrophy is associated with cancer-like metabolic reprogramming and limited by PHGDH

**DOI:** 10.1101/2020.12.01.403949

**Authors:** Lian E.M. Stadhouders, Sander A.J. Verbrugge, Jonathon A.B. Smith, Brendan M. Gabriel, Tim D. Hammersen, Detmar Kolijn, Ilse S.P. Vogel, Abdalla D. Mohamed, Gerard M.J. de Wit, Carla Offringa, Willem M. Hoogaars, Sebastian Gehlert, Henning Wackerhage, Richard T. Jaspers

## Abstract

Muscle fiber size and oxidative metabolism are inversely related, suggesting that a glycolytic metabolism may offer a growth advantage in muscle fibers. However, the mechanisms underlying this advantage remains unknown. Nearly 100 years ago, Warburg reported that cancer cells take up more glucose to produce glycolytic intermediates for anabolic reactions such as amino acid-protein synthesis. The aim of this study was to test whether glycolysis contributes to anabolic signalling responses and hypertrophy in post-mitotic muscle cells. Skeletal muscle hypertrophy was induced in vitro by treating mouse C2C12 myotubes with IGF-1. ^14^C glucose was added to differentiation medium and radioactivity in isolated protein was measured. We exposed differentiated C2C12 and primary mouse myotubes, to 2-deoxyglucose (2DG) and PHGDH siRNA upon which we assessed myotube diameter and signaling pathways involved in the regulation of muscle fiber size. Here, we present evidence that, hypertrophying C2C12 myotubes undergo a cancer-like metabolic reprogramming. First, IGF-1-induced C2C12 myotube hypertrophy increases shunting of carbon from glucose into protein. Second, reduction of glycolysis through 2-deoxy-D-glucose (2DG) lowers C2C12 and primary myotube size 16-40%. Third, reducing the cancer metabolism-associated enzyme PHGDH decreases C2C12 and primary myotube size 25-52%, whereas PHGDH overexpression increases C2C12 myotube size ≈20%. Fourth, the muscle hypertrophy-promoting kinase AKT regulates PHGDH expression. Together these results suggest that glycolysis is important for hypertrophying C2C12 myotubes by reprograming their metabolism similar to cancer cells.

## 1. Introduction

Having sufficient muscle mass and strength is associated with low morbidity and mortality(Gabriel and Zierath, 2017; Wolfe, 2006). An individual’s muscle mass and strength depend both on genetics(Arden and Spector, 1997; Verbrugge et al., 2018) and on environmental factors. Of the environmental factors, resistance (strength) training increases muscle mass and strength in most individuals.(Ahtiainen et al., 2016) Resistance training increases muscle mass by elevating protein synthesis for up to 48 h(McGlory et al., 2017) or even 72 h post-exercise (Miller et al., 2005), resulting in a positive protein balance in fed individuals.

The main mechanism by which resistance exercise increases protein synthesis is the activation of the serine/threonine kinase mTOR which is part of the mTORC1 complex (Goodman, 2019). In addition, hypertrophy-inducing stimuli such as synergist ablation (Chaillou et al., 2013) and acute resistance exercise (Pillon et al., 2020; Vissing and Schjerling, 2014) extensively change gene expression. Here, one of the most robust changes is the increased expression of the transcription factor Myc, whose expression increases >10-fold in synergist-ablated, hypertrophying mouse plantaris (Chaillou et al., 2013) and ≈6-fold 2.5 h after resistance exercise in human vastus lateralis muscle (Pillon et al., 2020; Vissing and Schjerling, 2014).

Both *MTOR* and *MYC* are cancer genes (Lawrence et al., 2013) and one of their functions is to contribute to the metabolic reprogramming seen in cancer cells (DeBerardinis and Chandel, 2016). Nearly 100 years ago, the metabolic reprograming of cancer cells was first experimentally demonstrated by Otto Warburg (Warburg et al., 1927). In a key experiment, the Warburg group compared glucose uptake and lactate production of sarcomas with that of healthy organs in rats. They noted that sarcomas took up more glucose and produced more lactate than other organs such as the liver, kidney or brain (Warburg et al., 1927). This demonstrated that cancer cells take up more glucose and have a higher glycolytic flux in the presence of oxygen. This phenomenon was termed *Warburg effect* by Efraim Racker in contrast to anaerobic glycolysis or the *Pasteur effect* (Racker, 1972). The purpose of the metabolic reprogramming of cancer cells was long poorly understood. Today we know that the pathways affected by metabolic reprogramming vary greatly between different types of cancer (Gaude and Frezza, 2016), both glycolysis and oxidative phosphorylation can be upregulated (DeBerardinis and Chandel, 2016) and that a key function of this metabolic reprogramming is to shunt glycolytic intermediates and other metabolites into anabolic reactions. These anabolic reactions include amino acids into protein and nucleotides into RNA/DNA synthesis, and help cancer cells to produce the biomass necessary for growth and proliferation (DeBerardinis and Chandel, 2016).

Given that the metabolic reprograming regulators mTORC1 and MYC are also active in a hypertrophying muscle, the question arises: Do hypertrophying skeletal muscle fibers reprogram their metabolism in a similar way to that of cancer cells? Several lines of evidence seem to support this idea. First, resistance exercise not only increases protein synthesis (McGlory et al., 2017) but also glucose uptake for at least one day post-exercise (Fathinul and Lau, 2009; Marcus et al., 2013). Second, Semsarian et al. noted that the induction of C2C12 hypertrophy through IGF-1 was additionally associated with increased lactate synthesis and an elevated expression of lactate dehydrogenase (Semsarian et al., 1999), which is essentially the Warburg effect. Similarly, muscle activation of AKT1, a known cancer metabolism regulator (Elstrom et al., 2004), not only causes muscle hypertrophy, but also increases the expression of glycolytic enzymes (Izumiya et al., 2008). Similarly, mTORC1 activation through a loss of its inhibitor NPRL2, results in muscle hypertrophy and induces aerobic glycolysis in mice (Dutchak et al., 2018). Finally, a loss of myostatin both induces muscle hypertrophy and promotes a shift to a more glycolytic metabolism (Mouisel et al., 2014). Collectively, these data suggest that the stimulation of muscle hypertrophy - through increased IGF1-AKT1-mTORC1 or reduced myostatin signaling - is associated with increased glycolysis and a metabolic reprogramming reminiscent to that which occurs in cancer cells (DeBerardinis and Chandel, 2016).

A specific cancer metabolism-associated enzyme that may contribute to such metabolic reprogramming during muscle hypertrophy is 3-phosphoglycerate dehydrogenase (PHGDH, E.C. 1.1.1.95). PHGDH channels 3-phosphoglycerate out of glycolysis, into serine biosynthesis and one-carbon metabolism, which is essential for nucleotide and amino acid synthesis, epigenetics and redox defense (Ducker and Rabinowitz, 2017; Reid et al., 2018). The knockdown of *PHGDH* inhibits proliferation of certain cancers (Possemato et al., 2011), endothelial cells (Vandekeere et al., 2018), and fibroblasts (Hamano et al., 2018), suggesting that PHGDH-mediated metabolic reprograming is important for proliferation and cellular growth. Besides, muscle stem cells express more *Phgdh* when they become activated and start to proliferate (supplementary data of Ryall et al., 2015)). In addition, *Phgdh* mRNA expression also increases in terminally differentiated pig muscles when hypertrophy is stimulated with the β2-agonist ractopamine (Brown et al., 2016). Together, this shows that PHGDH becomes activated and/or more abundant in proliferating cells and in at least one type of skeletal muscle hypertrophy.

Currently, it is unclear whether a hypertrophying muscle reprograms its metabolism similar to cancer cells so that glycolytic intermediates and other metabolites are shunted out of energy metabolism into anabolic reactions such as serine biosynthesis and one-carbon metabolism. The aim of this study was therefore to investigate whether hypertrophying C2C12 myotubes shunt more carbon from glucose into amino acid and nucleotides for protein and RNA synthesis. We also investigated whether inhibition of glycolytic flux and *Phgdh* knockdown or overexpression affected C2C12 myotube size, in untreated or IGF-1-treated myotubes. We found that hypertrophying C2C12 and primary mouse myotubes indeed shunt more carbon from glucose into protein and RNA synthesis, that the inhibition of glycolysis and knockdown of *Phgdh* reduces myotube size whereas overexpression of *Phgdh* increases myotube size, respectively. Collectively, this suggests that glycolysis is important for hypertrophying C2C12 myotubes which reprogram their metabolism similar to cancer.

## 2. Materials and methods

### 2.1 C2C12 cell culture

C2C12 muscle cells (ATCC, Cat# CRL-1772, RRID:CVCL_0188; Middlesex, UK; cells are regularly tested for contamination) were grown to confluency in growth medium containing Dulbecco’s Modified Eagle’s Medium DMEM (Gibco, Cat#31885, Waltham, MA, USA), containing 10% fetal bovine serum (Biowest, Cat#S181B, Nuaillé, France), 1% penicillin/streptomycin (Gibco, Cat#15140, Waltham, MA, USA) and 0.5% amphotericin B (Gibco, Cat#15290-026, Waltham, MA, USA) and incubated at 37°C in humidified air with 5% CO_2_. Once 90% confluent, medium was changed to differentiation medium consisting of DMEM supplemented with 2% horse serum (HyClone, Cat#10407223, Marlborough, MA, USA) and 1% penicillin/streptomycin. This medium was refreshed daily for 3 days until treatment.

### 2.2 Primary myoblast culture

Primary muscle stem cells where obtained from extensor digitorum longus (EDL) muscles of 6-week to 4-month old mice of a C57BL/6 background. The experiments were conducted with post-mortem material from surplus mice (C57BL/6J) originating from breeding excess that had to be terminated in the animal facility, and therefore, these animals do not fall under the Netherlands Law on Animal Research in agreement with the Directive 2010/63/EU. The EDL muscles were incubated in collagenase type I (Sigma-Aldrich, Cat#C0130, Saint Louis, MO, USA) at 37 °C, in air with 5% CO_2_ for 2 h. The muscles were washed in DMEM, containing 1% penicillin/streptomycin (Gibco, Cat#15140, Waltham, MA, USA) and incubated in 5% Bovine serum albumin (BSA)-coated dishes containing DMEM for 30 min at 37 °C in air with 5% CO_2_ to inactivate collagenase. Single muscle fibres were separated by gently blowing with a blunt ended sterilized Pasteur pipette. Subsequently, muscle fibres were seeded in a thin layer matrigel (VWR, Cat#734-0269, Radnor, PA, USA)-coated 6-well plate containing DMEM growth medium, 1% penicillin/streptomycin (Gibco, 15140, Waltham, MA, USA), 10% horse serum (HyClone, Cat#10407223, Marlborough, MA, USA), 30% fetal bovine serum (Biowest, Cat#S181B, Nuaillé, France), 2.5ng ml-1 recombinant human fibroblast growth factor (rhFGF) (Promega, Cat#G5071, Madison, WI, USA), and 1% chicken embryonic extract (Seralab, Cat#CE-650-J, Huissen, The Netherlands). Primary myoblasts were allowed to proliferate and migrate off the muscle fibres for 3-4 days at 37 °C in air with 5% CO_2_. After gentle removal of the muscle fibres, myoblasts were cultured in matrigel-coated flasks until passage 5. Cells were pre-plated in an uncoated flask for 15 min with each passage to reduce the number of fibroblasts in culture. Cell population was 99% Pax7^+^. All experiments with primary myoblasts were performed on matrigel-coated plates. Primary myoblasts were cultured in differentiation medium to differentiate for 2 days until treatment.

### 2.3 Cell treatments

#### 2.3.1 ^14^C-glucose to protein/RNA flux analysis

After 48 h in differentiation medium, myotubes were incubated in differentiation medium plus treatment for up to 48 h. Treatments were one of: vehicle control (0.00001% bovine serum albumin; BSA), IGF-1 (100 ng ml-1; recombinant Human IGF-1, Peprotech, Cat#100-11, London, UK, rapamycin (100 ng ml-1; Calbiochem, Cat#553210, Watford, Hertfordshire, UK) and IGF-1 + rapamycin.

#### 2.3.2 Inhibition of glycolysis via 2-deoxyglucose (2DG)

Differentiated C2C12 myotubes were treated with IGF-1, 2DG or both. IGF-1 (100 ng ml-1) and 2DG (5 mM, Cat#D6134, Sigma Aldrich) were diluted in differentiation medium. On day 3, IGF-1 and 2DG were added to the myotube culture for 24 hours. For inhibition of AKT, we used the compound MK-2206 (10-1000 μM, Bio-Connect, Cat#HY-10358, Netherlands). Differentiation medium was refreshed daily for 4 days. On day 7, myotubes were treated with MK-2206 (1000 μM). After 1 h incubation with MK-2206 (10 μM, 100 μM or 1000 μM), IGF-1 (100 ng ml-1) was added. Cells were harvested on day 8, after 24 h of IGF-1 treatment.

### 2.4 Protein determination

Cells were lysed on ice in 500 μl of radioimmunoprecipitation assay (RIPA) buffer (Sigma-Aldrich, R0278, Saint Louis, MO, USA) supplemented with phosphatase (1:250, Sigma-Aldrich, 04906837001, Saint Louis, MO, USA) and proteinase (1:50, Sigma-Aldrich, Cat#11836153001, Saint Louis, MO, USA) inhibitor cocktails, 0.5 M Ethylenediaminetetraacetic acid (EDTA; 1:500), sodium fluoride (NaF; 1:50) and sodium orthovanadate (1:50). Lysates were left on ice for 15 min and cellular debris removed by centrifuging at 13,000 rpm for 15 min, at 4°C. Supernatants were then transferred into fresh Eppendorfs and frozen at −80°C (for radiolabeled glucose to protein) or their protein concentrations calculated (Western blot) using a Pierce BCA Protein Assay kit (Thermo Scientific, Cat#23225, Waltham, MA, USA).

### 2.5 RNA isolation

After washing cells with Phosphate-Buffered Saline (PBS). Cells were lysed in TRI reagent (Invitrogen, 11312940, Carlsbad, CA, USA) and stored at −80°C. RNA was isolated using RiboPureTMkit (Applied Biosystems, Foster City, CA, USA) and converted to cDNA with high-capacity RNA to cDNA master mix (Applied Biosystems, Foster City, CA, USA). cDNA was diluted 10x and stored at −20°C.

### 2.6 ^14^C-glucose to protein flux analysis

1 μl ml-1 of 0.1 mCi ml-1 ^14^C glucose (PerkinElmer, Cat# NEC042V250UC, Waltham, MA, USA) was added to differentiation medium 48 h before the end of the experiments. 200 μl of medium was maintained and added to 4 ml of scintillation fluid to acquire initial radioactivity readings for rate of incorporation calculations. To separate protein from other macromolecules, harvested lysates of 48 h treated cells were fractionated by acetone precipitation. The protein pellet was then re-suspended in PBS and incubated in 4 mL of scintillation fluid (Insta-Gel Plus, PerkinElmer) for 24 h (to homogenize the samples) before measuring the radioactivity in a scintillation counter. Results are given in counts per minute (CPM) per 10 cm diameter Petri dish.

### 2.7 Gel-phosphorimaging

Protein was extracted as described in the section “Protein determination”. Samples were prepared and electrophoresed as described in the section ‘Western Blotting’. The 12% Criterion XT pre-cast Bis-Tris gel (Bio-rad, Cat# 3450119, Hemel Hempstead, UK) was then stained with Silver Stain (Bio-rad, Cat#1610481, Hemel Hempstead, UK) and imaged as a loading control quality check. The gel was then dried using a gel dryer and incubated with an imaging plate inside a radiography cassette. After 48 h the imaging plate was imaged using a phosphor-imager (Fuji FLA3000).

### 2.8 ^14^C-glucose to RNA flux analysis

After the 48 h incubation with 1 μl ml-1 of 0.1 mCi ml-1 ^14^C glucose and reagents, RNA was extracted using an RNeasy Mini Kit (Qiagen, Cat#74104, Valencia, CA, USA) according to the manufacturer’s instructions. The elution was then placed in 4 ml of scintillation fluid for 24 h prior to measuring radioactivity of the samples in a scintillation counter.

### 2.9 Western blotting

Respective volumes of lysate were diluted in 5 times Laemmli SDS buffer and denatured for 5 min at 95°C, prior to western blotting. Samples were then electrophoresed on 12% Bis-Tris gels (Bio-rad, Cat#3450125, Hemel Hempstead, UK) and transferred onto PVDF membranes (GE Healthcare, Cat#15269894, Chicago, IL, USA) using a semi-dry transfer blotter (Bio-rad). Membranes were blocked in prime blocking agent (GE Healthcare, Cat#RPN418, Chicago, IL, USA), then incubated with primary Phospho-P70S6K (Thr389; 1:2000; Cell Signaling Technology, Cat# 9234, Leiden, The Netherlands, RRID:AB_2269803), Phospho-AKT Ser473 (1:2000; Cell Signaling Technology, Cat# 4060, Leiden, The Netherlands, RRID:AB_2315049), α-TUBULIN (1:10000; Cell Signaling Technology, Cat# 2125, Leiden, The Netherlands, RRID:AB_2619646), pan-ACTIN (1:1000, Cell Signaling Technology, Cat# 8456, Leiden, The Netherlands, RRID:AB_10998774), PHGDH (1:1000; Cell Signaling Technology, Cat# 13428, Leiden, The Netherlands, RRID:AB_2750870), Phospho-AMPK (Thr172, 1:500, Cell Signaling Technology, Cat# 2531, Leiden, The Netherlands, RRID:AB_330330), and anti-rabbit/mouse IgG secondary antibody (1:2000; Roche, Cat#12015218001, Basal, Switzerland) prior to fluorescent imaging. Densities of the bands from blot images were normalized loading control by densitometric analysis using ImageJ software (RRID:SCR_003070).

### 2.10 Myotube size measurement

Four photographs of each well were taken at 10x magnification after the 24 h treatment. Diameters were measured in 20-50 myotubes at 5 equidistant locations along the length of the cell using ImageJ (http://rsbweb.nih.gov/ij/, National Institutes of Health, Bethesda, MD, USA; RRID:SCR_003070) and taking into account the pixel-to-aspect ratio.

### 2.11 Lactate concentration

Lactate levels in the culture medium were measured using Lactate Assay Kit (Sigma-Aldrich, Cat#MAK064, Saint Louis, MO, USA) according to manufacturer’s instructions. Culture medium samples were directly deproteinized with a 10 kDA MWCO spin filter to ensure lactate dehydrogenase was separated from the medium. Samples (0.5 μL) were assayed in duplicate on a 96 well plate. Lactate Assay Buffer was added to bring samples to a final volume of 50 μL/well. Lactate levels were determined by colorimetric assay on 570 nm and concentrations were based on the standard curve.

### 2.12 Real-time quantitative PCR

cDNA was analysed using real-time quantitative PCR (see Table 1 for primer details). Experiments were conducted in duplicates. Concentration of the transcriptional target was detected with fluorescent SYBR Green Master Mix (Fischer Scientific, Cat#10556555, Pittsburgh, PA, USA). Transcriptional expressions of the target genes were referenced to 18S housekeeping gene. Relative changes in gene expression were determined with the ΔCt method.

**Table 1.**
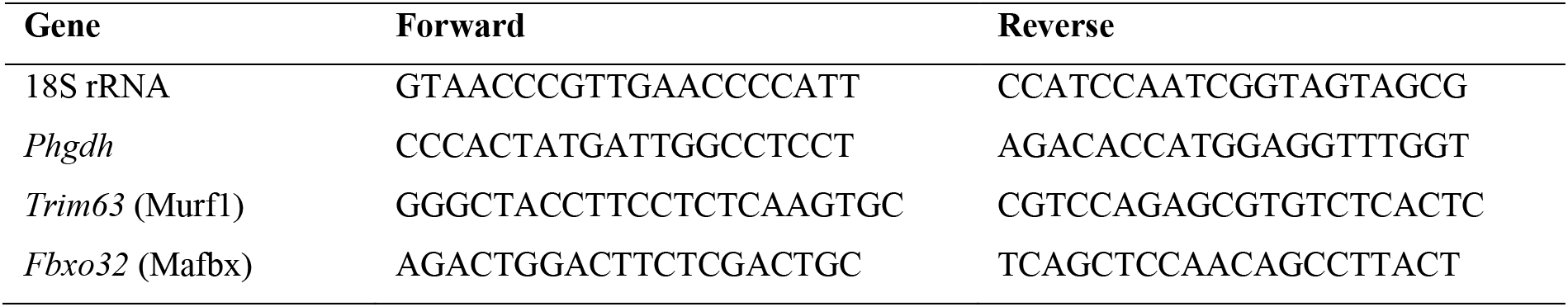
PCR primers

### 2.13 siRNA-mediated knockdown of *Phgdh*

To carry out a PHGDH loss-of-function experiment, we knocked down *Phgdh* in C2C12 and primary myotubes using silencer RNA (Ambion, Carlsbad, CA, USA, see Table 2 for siRNA sequences). C2C12 myoblasts or primary myoblasts were grown and differentiated as described. On day 6, myotubes were transfected with siRNA targeted against *Phgdh* using the liposome-mediated method (Lipofectamine RNAiMAX, Invitrogen, Cat# 13778100, Carlsbad, CA, USA). As a negative control, a non-targeting silence RNA sequence (siControl) was used. siRNA was diluted in Opti-MEM medium and incubated for 5-10 minutes with Lipofectamine mixture. RNA-lipofectamine complexes with a final concentration of 20 nM were added to each well. On day 7, the differentiated myotubes were treated with IGF-1 (100 ng ml-1) and harvested at day 8 (48 hours post-transfection).

**Table 2.**
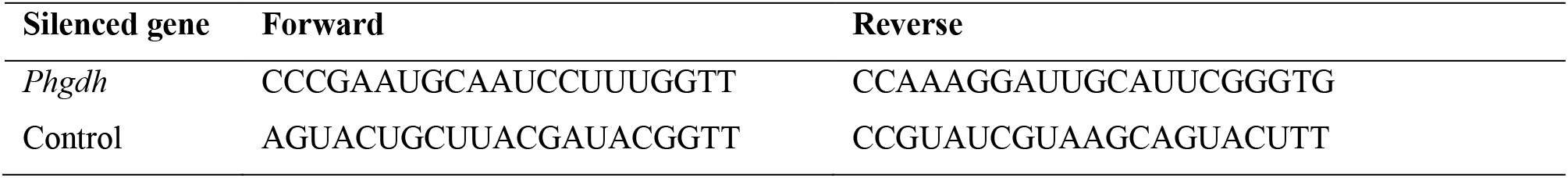
siRNA information

### 2.14 PHGDH plasmid cloning, retrovirus and retroviral infection

To carry out a PHGDH gain-of-function experiment, we subcloned a human PHGDH pMSCV retroviral vector and transduced C2C12 myoblasts with this vector prior to differentiation. Human PHGDH cDNA (transcript variant NM_006623.4, which encodes a protein of 533 amino acids) was amplified by RT-PCR and cloned into pMSCV-IRES-eGFP using In-Fusion® HD Cloning Kit User Manual (Takara, Cat#638920, Shiga, Japan) following the manufacturer’s instructions (see Table 3 for primer sequences).

**Table 3.**
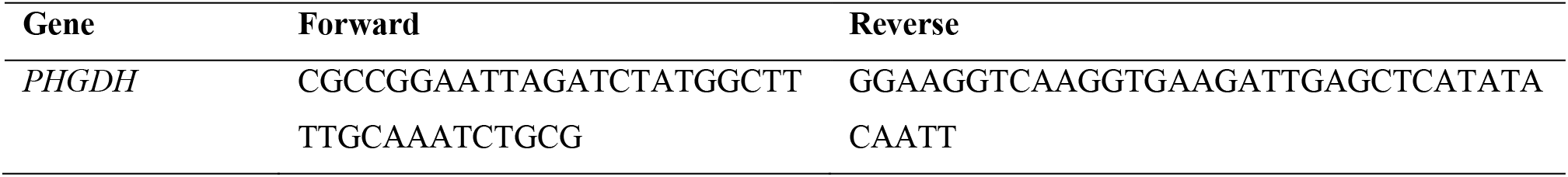
Primers

For retroviral particle production, HEK293T cells were seeded at a density of 3 × 10^6^ cells per T75 flask, 24 h prior to transfection. 1 h before transfection, medium was changed to 7 ml of fresh growth medium, DMEM Glutamax (Gibco, Cat#10566016, Waltham, MA, USA) + 10% FBS (Biowest, Cat#S181B, Nuaillé, France). For transfection, 4 μg of PHGDH plasmid or 4 μg of empty vector plasmid (Addgene, plasmid #52107) was mixed with 4 μg of DNA RV helper plasmid (Addgene, plasmid #12371), in 1800 μl of Opti-MEM reduced serum medium (ThermoFisher, Cat# 31985070, Waltham, MA, USA). 6 μl of Lipofectamine Plus reagent was then added and incubated for 5 min at room temperature, followed by 24 μl of Lipofectamine LTX (ThermoFisher, Cat#15338100, Waltham, MA, USA) for a further 30 min incubation at room temperature. Medium was changed 6 and 24 h after transfection, 48 h post transfection, medium was changed to 7 ml of fresh growth medium. Retroviral particles were collected 12, 24 and 36 h after the last medium change by collecting all 7 ml of medium, and filtrating through 0.45 μm filters.

For retroviral infection, 3 × 10^4^ C2C12 cells were seeded in a 6-well plate overnight at 37°C in a 5% CO_2_ incubator. 1 h prior to infection, medium was changed to 1.5 ml of fresh growth medium and cells were infected by adding retroviral particles in a ratio of 1:4. Cells were incubated until reaching 90% confluence and then differentiated as described.

### 2.15 Statistical analysis

Shapiro–Wilk tests were used to test for normal distribution. Data were then analysed using unpaired t-test, two-way analysis of variance (ANOVA), or three-way ANOVA for normally distributed data. When data was not normally distributed, we used the Mann-Whitney U test. In the case of a significant ANOVA effect, a Bonferroni test was used to determine significant differences between conditions. Significance was set at *p*<0.05. Data are presented as mean ± SEM with individual data points. Statistical analyses were performed using Prism 7.0 (GraphPad Prism, RRID:SCR_002798).

## 3 Results

### 3.1 IGF-1 induced myotube growth increases 14C-incorporation into protein and RNA

A key feature of metabolic reprogramming in cancer is that glycolytic intermediates and other metabolites are shunted out of energy metabolism and into anabolic pathways (DeBerardinis and Chandel, 2016). In relation to skeletal muscle hypertrophy, a key question is: does a similar shunting of metabolites happen in hypertrophying muscles? To try to answer this question, we stimulated hypertrophy of C2C12 myotubes with 100 ng ml-1 of IGF-1 (Rommel et al., 2001) and measured the rate by which 14C derived from 14C-glucose is incorporated into amino acid→protein and nucleotide→RNA synthesis (Figure 1a). This experiment revealed that 14C-incorporation both into protein and RNA was already measurable at baseline and that IGF-1 increased 14C-incorporation into protein by ≈71%, on average, versus control (12924 ± 2113 versus 7581 ± 1586 CPM) (Figure 1c). Generally, little 14C was incorporated into RNA and whilst IGF-1 increased 14C incorporation into RNA (two-way ANOVA, *p*=0.030), this was only a non-significant trend (Bonferonni post-hoc) (Figure 1d). Additional treatment with the mTOR-inhibitor rapamycin reduced IGF-1-stimulated 14C incorporation into protein by ≈61% when compared to control, and ≈77% when compared to IGF-1 stimulation, which suggests that the 14C incorporation into protein is mTORC1 dependent. Together this data indicates that C2C12 myotube hypertrophy is associated with an increased shunting of carbon from glucose into anabolic reactions, such as incorporation of amino acid into proteins.

**Figure 1.**
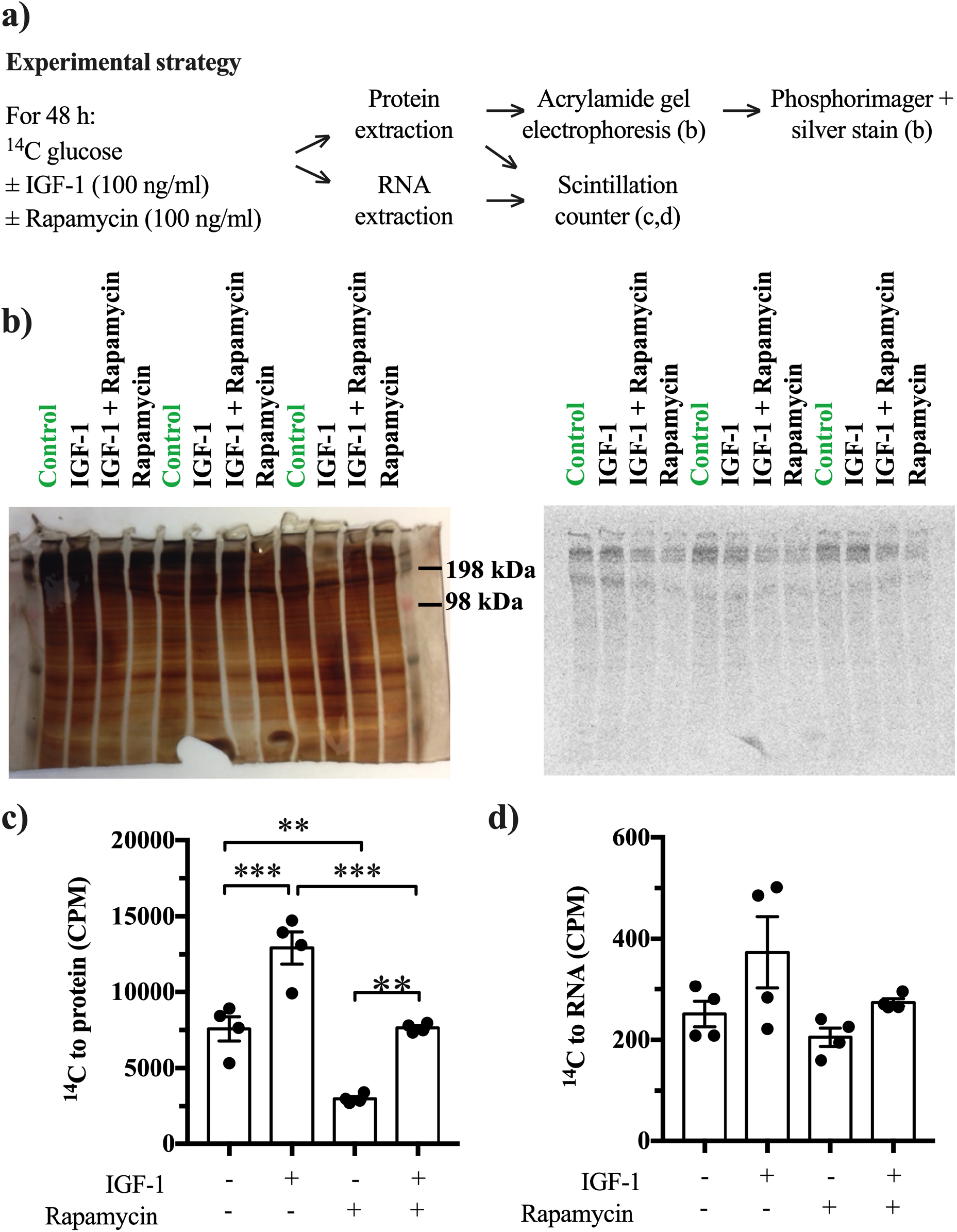
Glucose-derived (14C) carbon can be converted into protein and RNA particularly in IGF-1 hypertrophy stimulated myotubes, suggesting cancer-like metabolic reprogramming so that energy metabolites are channeled into anabolic pathways. (a) Schematic depiction of the experimental strategy. (b) Dried gels showing all protein bands and radioactivity detected in the gel using a phospho-imager (n=3 samples per treatment on one gel). (c,d) Quantification of radioactivity (n=4 samples per treatment; in counts per minute; CPM) per 10 cm diameter dish in (c) precipitated and isolated protein lysates and (d) extracted RNA (n=4). *Significantly different between indicated conditions, two-way ANOVA with Bonferonni post-hoc test (*p*<0.05).

### 3.2 Blocking glycolysis inhibits myotube growth

Because glycolysis is not only a key metabolic pathway but also a feeder pathway for anabolic reactions (DeBerardinis and Chandel, 2016), we tested whether an inhibition of glycolysis affected C2C12 myotube size and hypertrophy. For this purpose, we inhibited glycolysis with 2-deoxy-D-glucose (2DG; Xi et al., 2014) and then measured the diameter of control myotubes and myotubes stimulated with IGF-1. We found that 2DG treatment reduced lactate concentrations as expected (Figure 2a) and reduced C2C12 myotube diameter by ≈30%, on average, in untreated myotubes and by ≈40% in IGF-1-treated myotubes (Figure 2b,c). Also in primary muscle cells, myotube size decreased after 2DG exposure (Figure 2d). Collectively, these data suggest that inhibition of glycolytic flux reduces myotube size.

**Figure 2.**
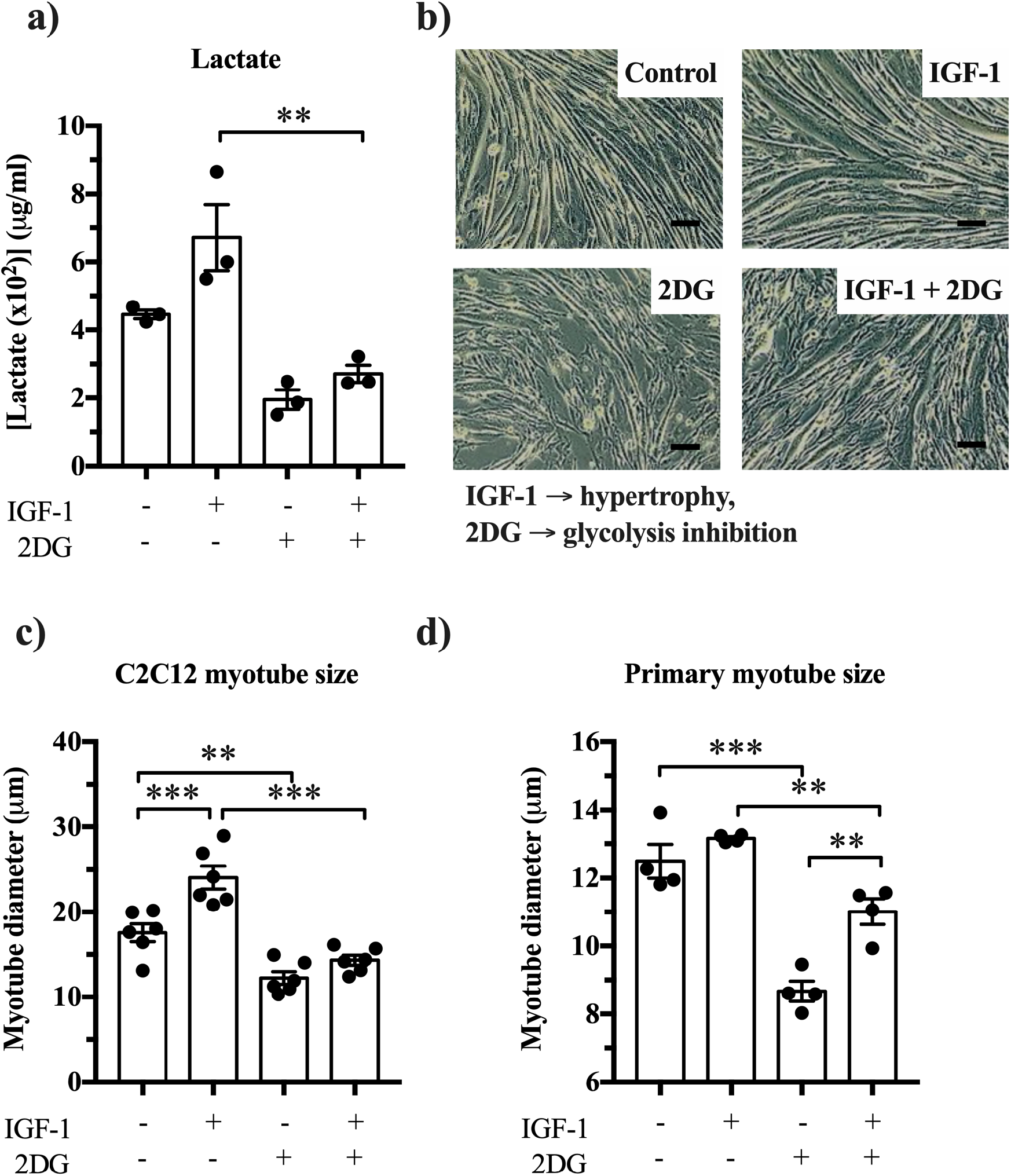
Glycolysis inhibition reduces myotube diamter in C2C12 and primary muscle cells. (a) Effects of 2DG on lactate concentrations (n=3), (b,c) C2C12 myotube diameter (n=5) and (d) primary myotubes (n=4). Scale bar is 100 μm. *Significantly different between indicated conditions, unpaired t-test, Mann-Whitney U or two-way ANOVA with Bonferonni post-hoc test (*p*<0.05).

### 3.3 Inhibition of glycolysis and IGF-1 affect protein turnover and PHGDH

The inhibition of glycolytic flux through 2DG may not only affect the generation of glycolytic intermediates as substrates for anabolic reactions but also energy-sensitive signalling mechanisms. To answer this, we measured activity markers and the expression, phopho-AMPK and atrophy-associated E3 ubiquitin ligases. We observed no effect of 2DG on AMPK phosphorylation (Figure 3a; two-way ANOVA, *p=0.808),* indicating that any potential diminished energy state did not cause the reduced myotube size. On the other hand, we found that 2DG decreases P70S6K phosphorylation (Figure 3b), repressing protein synthesis. While *Trim63* remains unaffected by blocking glycolysis (Figure 3e), the other protein degradation marker, *Fbxo32,* is attenuated by 2DG (Figure 3d). Together this shows that 2DG-associated inhibition of myotube hypertrophy stems from the suppression of both protein synthesis and protein degradation.

**Figure 3.**
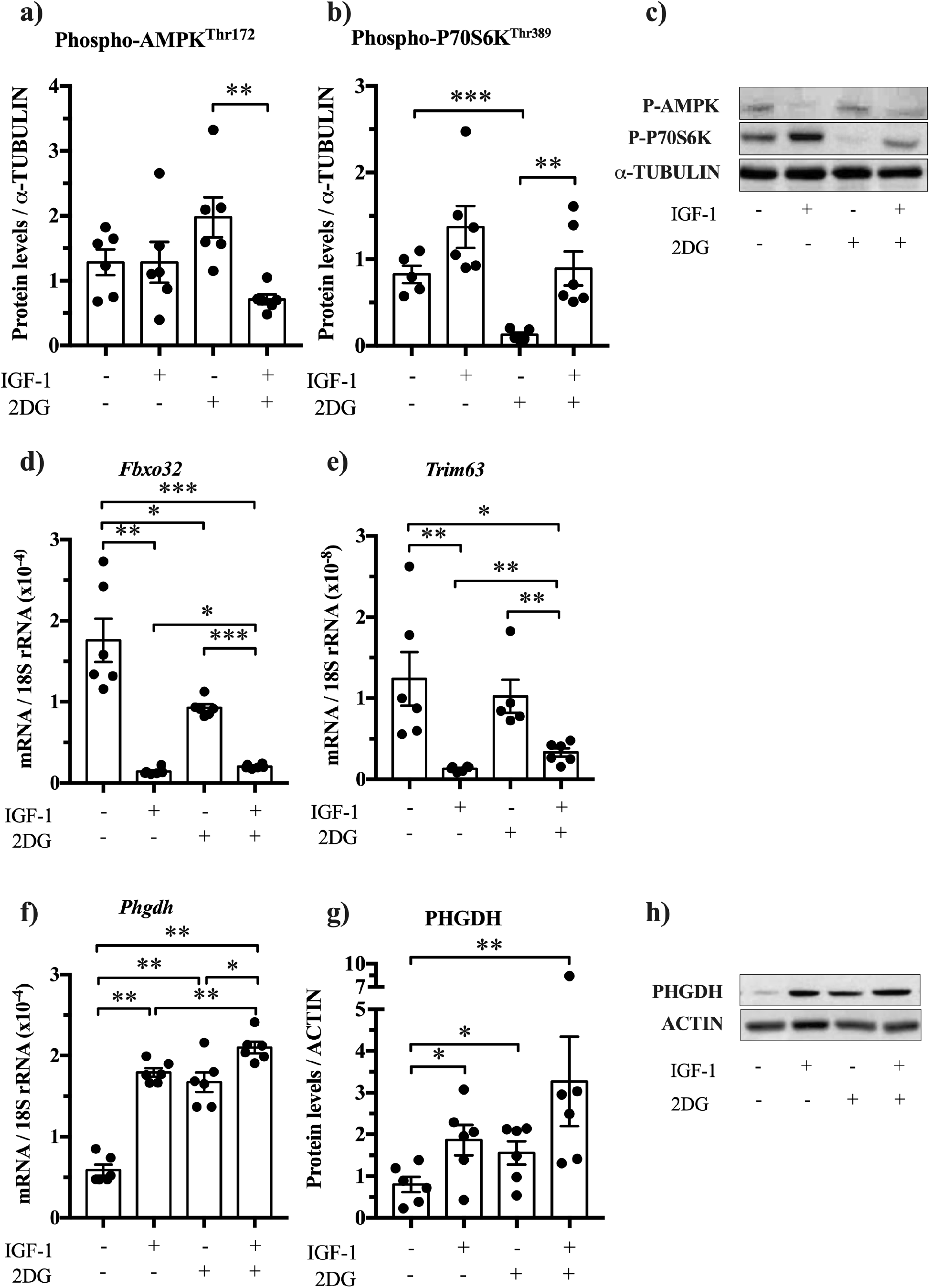
2DG affects protein abundance and mRNA expression in myotubes (a) No effect of 2DG on Phospho-AMPK, but an interaction effect was observed (p=0.018) indicating that IGF-1 hampers 2DG-induced AMPK phosphorylation. (b) P70S6K phosphorylation at residue Thr389 increases upon IGF-1 stimulation (p=0.002) and decreases after 2DG exposore (p=0.004). (c) Western blots for Phospho-AMPK and Phospho-P70S6K with α-ACTIN as loading control. (d) Mafbx (*Fbxo32*) and (e) Murf1 (*Trim63*) decrease upon IGF-1 stimulation (p=0.010 and p<0.001, respectively). (f) IGF-1 and 2DG increase *Phgdh* mRNA expression and (g,h) PHGDH abundance in C2C12 (n=6). *Significantly different between indicated conditions, unpaired t-test, Mann-Whitney U test or two-way ANOVA with Bonferonni post-hoc test (*p*<0.05).

### 3.4 Knock-down of PHGDH attenuates myotube growth

Next we studied the role of the cancer reprogramming-associated enzyme PHGDH. PHGDH catalyses the first reaction of the serine biosynthesis pathway (3-phosphoglycerate + NAD+ ↔ 3-phosphooxypyruvate + H+ + NADH) and is important for one-carbon metabolism (Ducker and Rabinowitz, 2017). PHGDH is well-known to limit cell proliferation (Possemato et al., 2011), but also in post-mitotic growth it plays a role. Indeed, PHGDH and other serine biosynthesis enzymes are increased when stimulating muscle hypertrophy in pigs with the β2-agonist ractopamine (Brown et al., 2016). In agreement, we observed that C2C12 myotubes increase *Phgdh* expression by 204% upon IGF-1 stimulation, and in primary myotubes by 104% (Figure 3f,g). To investigate whether normal levels of PHGDH limit myotube size and hypertrophy, we knocked down PHGDH through siRNA-mediated RNA interference (Figure 4a-c) and determined the effect on myotube diameter in C2C12 and primary muscle cells (Figure 4d-g). siRNA interference resulted in a reduction of both *Phgdh* mRNA, to 30-50% of baseline levels (Figure 4a,g), and protein, non-significantly to ≈16% of baseline levels (Figure 4b,c). This knockdown of PHGDH decreased C2C12 myotube size under both basal and IGF-1-stimulated conditions, on average, ≈29% and ≈52%, respectively (Figure 4d,e). Consistently, we observed also in primary-derived myotubes decreased size upon PHGDH knockdown by 25% and 36% in control and after IGF-1 stimulation, respectively (Figure 4g). Together these results show that a knockdown of PHGDH reduces myotube size, which further supports the idea that a cancer-like metabolic reprogramming occurs in hypertrophying skeletal muscle.

**Figure 4.**
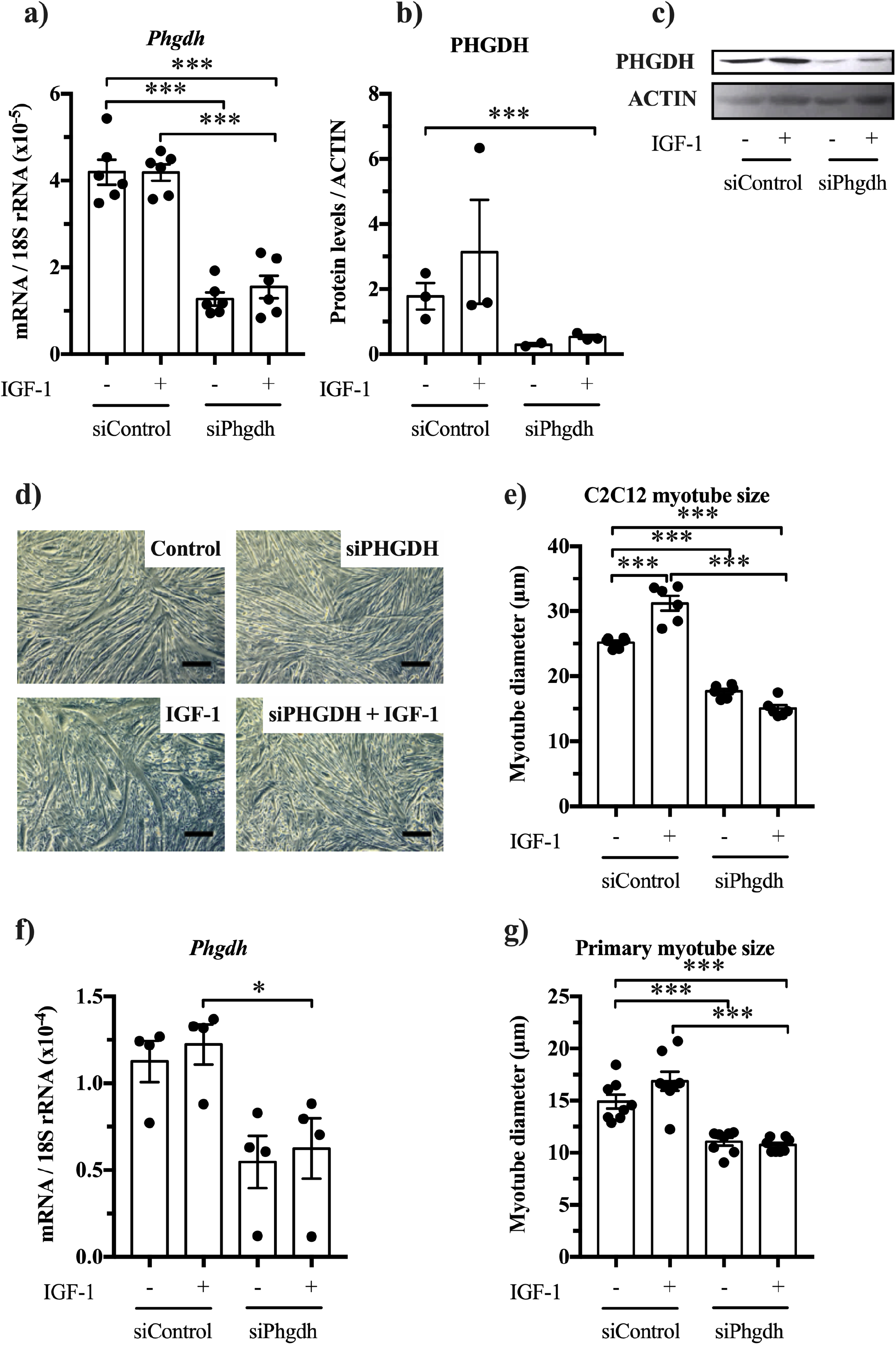
PHGDH knockdown by siRNA reduces C2C12 myotube size in control and IGF-1-stimulated myotubes. (a) *Phgdh* mRNA and (b,c) PHGDH protein levels after siRNA treatment. (d) Morphology of C2C12 myotubes, (e) the effect of PHGDH knockdown on C2C12 myotube diameter (n=6). (f,g) siRNA in primary myotubes decreased *Phgdh* mRNA and myotube size (n=4-6). Scale bar is 100 μm. *Significantly different between indicated conditions, unpaired t-test, Mann-Whitney U or two-way ANOVA with Bonferonni post-hoc test (*p*<0.05).

### 3.5 PHGDH overexpression increases myotube size

Because a loss of PHGDH reduced myotube size, we next investigated whether a gain of PHGDH would increase C2C12 myotube size. PHGDH mRNA expression increased after retroviral transduction (Figure 5a) and increased C2C12 myotube size by ≈20% independent of IGF-1-stimulation (Figure 5b,c). Furthermore, we observed a trend towards increased myotube size in untreated control cells after PHGDH overexpression compared to empty vector, which we also saw at 24 h in myotubes that were cultured in absence of IGF-1 (Figure 5b,c).

**Figure 5.**
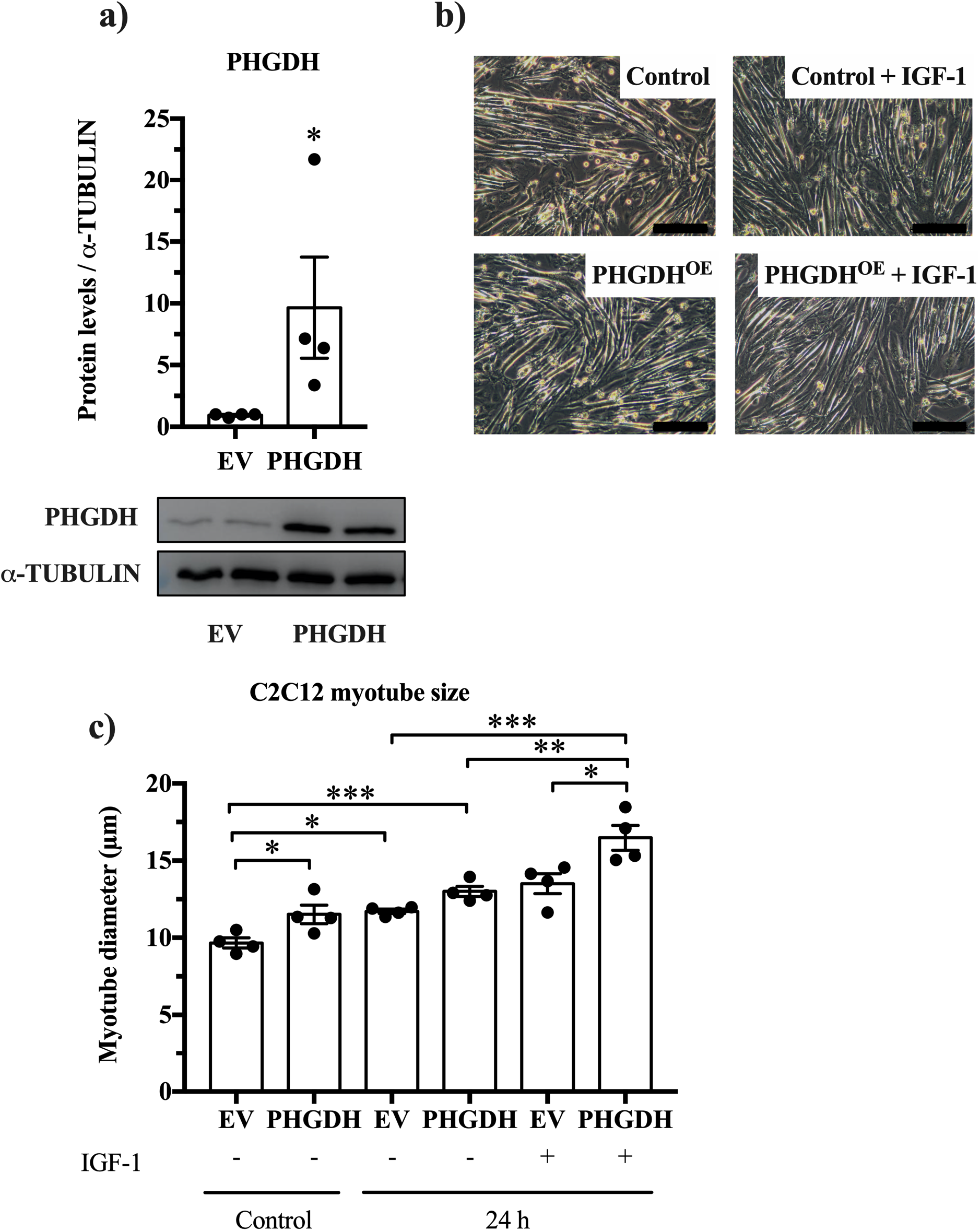
Overexpression of PHGDH (PHGDH^OE^) significantly increases PHGDH protein levels and myotube size (a) Overexpression of human PHGDH in C2C12 myotubes (n=4). Note that we could not distinguish between endogenous mouse PHGDH and overexpressed human PHGDH. (b,c) C2C12 myotube size increases upon PHGDH overexpression compared to empty vector (n=4). Scalebar is 200 μm. *Significantly different between indicated conditions, two-way ANOVA (overexpression x time and overexpression x IGF-1) with Bonferonni post-hoc test (*p*<0.05).

### 3.6 AKT regulates PHGDH in C2C12 myotubes

Activation of AKT stimulates aerobic glycolysis, i.e. the Warburg effect, in cancer cells (Elstrom et al., 2004), promotes muscle hypertrophy (Lai et al., 2004) and is associated with a shift towards glycolysis (Izumiya et al., 2008). We therefore wanted to assess whether AKT also regulates PHGDH e.g. through a change in PHGDH protein abundance. To study this, we used IGF-1 to activate AKT and stimulate C2C12 hypertrophy, and the AKT inhibitor, MK-2206, to repress AKT activity and measured Phgdh protein through western blotting. We confirmed the blocking effect of MK-2206 on activity-associated AKT (*p*=0.020) (Figure 6a) and P70S6K phosphorylation (Figure 6b), which was accompanied by reduced myotube diameter (Figure 6c,d). In addition, we found that in IGF-1-treated C2C12 myotubes, MK-2206 reduced PHGDH abundance in a dose dependent manner (by 42% at 1000 μM) (Figure 6e,f). This suggests that AKT regulates PHGDH which further supports the idea that a hypertrophying muscle reprograms its metabolism similar to cancer cells.

**Figure 6.**
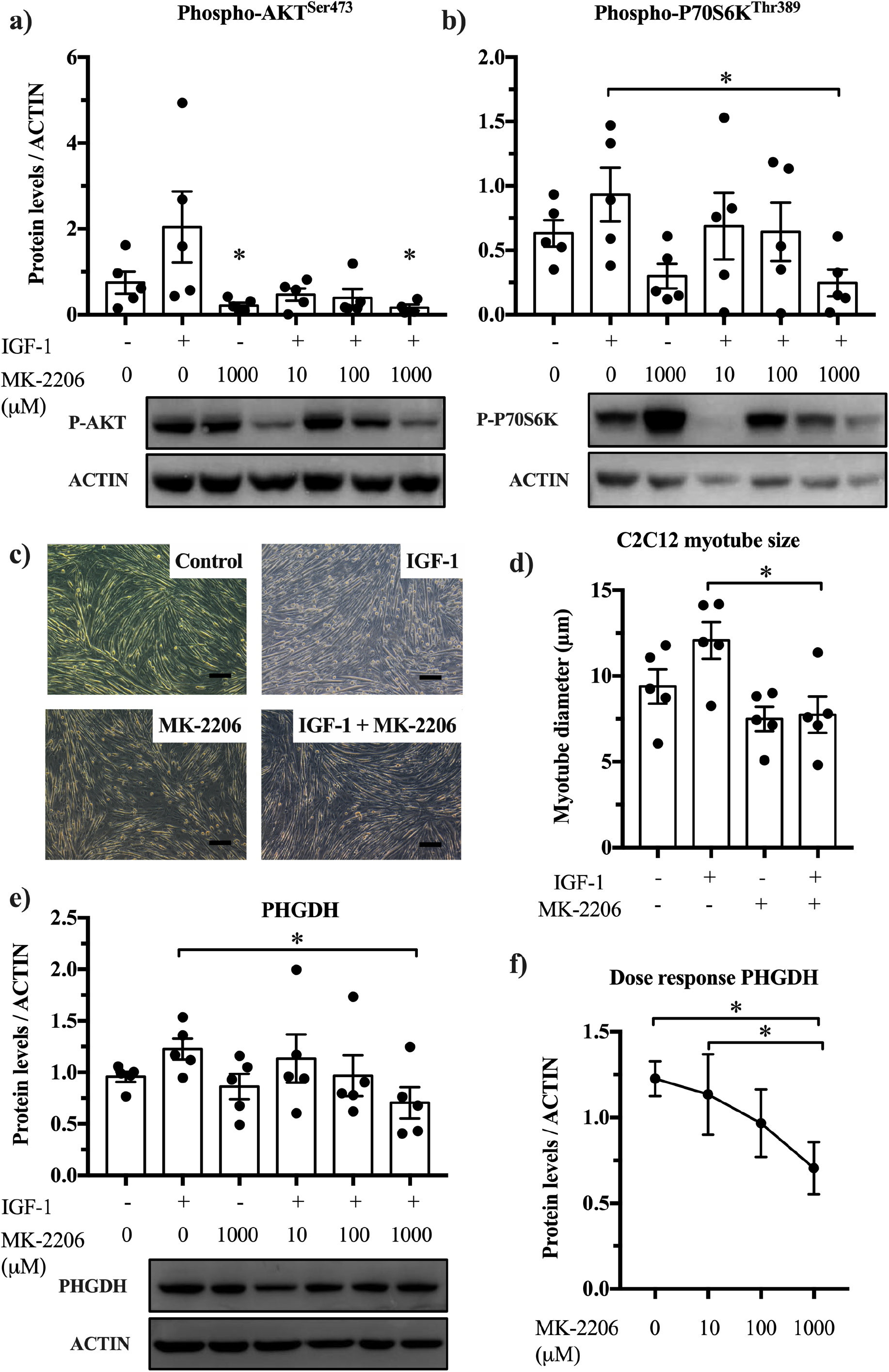
Blocking AKT prevents IGF-1 induced PHGDH upregulation (a,b) MK-2206 blocks AKT phosphorylation (two-way ANOVA, *p*=0.020) and P70S6K phosphorylation in untreated and IGF-1 treated myotubes (n=5). (c,d) The AKT blocker, MK-2206 (1000 μM), prevents IGF-1 induced hypertrophy in myotubes and decreases PHGDH abundance (e,f) in a dose-dependent manner (n=5; Friedman’s test). Scale bar is 100 μm. *Significantly different between indicated conditions, two-way ANOVA (IGF-1 x MK-2206, 1000 μM) with Bonferonni post-hoc test, or Friedman’s test (*p*<0.05).

## 4 Discussion

This study reports five findings that support the idea that hypertrophying muscles reprogram their metabolism similar to cancer (Figure 7). First, the stimulation of C2C12 myotube hypertrophy through IGF-1 increases shunting of carbon from glucose especially into amino acid→protein synthesis. Second, we confirm that IGF-1 increases glycolytic flux in C2C12 and primary myotubes (Semsarian et al., 1999) and report that a reduction of glycolytic flux through 2DG-mediated inhibition lowers myotube size. Third, a reduction of the serine biosynthesis-catalysing enzyme PHGDH decreases C2C12 and primary myotube size, suggesting that PHGDH limits normal myotube size and myotube hypertrophy. Fourth, the overexpression of PHGDH in C2C12 myotubes increases myotube diameter. Fifth, the muscle hypertrophy-promoting kinase AKT regulates the expression of PHGDH.

**Figure 7.**
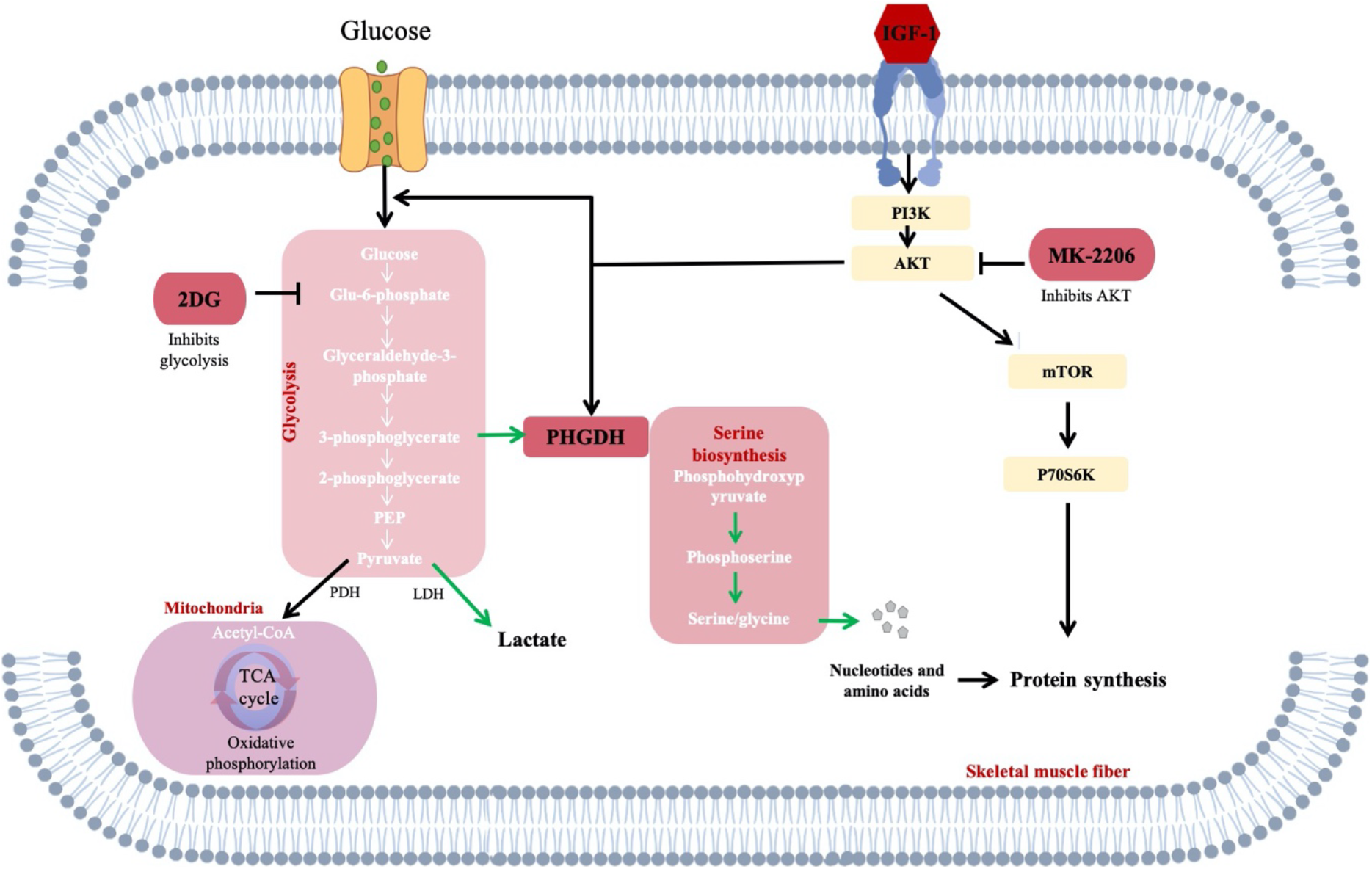
Glucose is taken up by muscle cells and via several glycolytic intermediates catalyzed to pyruvate for energy production in mitochondria. Alternatively, cancer-like metabolic remodeling shunts the glycolytic intermediate 3-phosphoglycerate (3PG) into *de novo* serine synthesis pathway. Phosphoglycerate dehydrogenase (PHGDH) catalyses the first step of 3PG to 3-phosphohydroxypyruvate which is then converted into 3-phosphoserine and ultimately serine. Serine is subsequently used for protein synthesis, or as a precursor for nucleotide synthesis. IGF-1 induces hypertrophy in post-mitotic muscle cells, which is abolished by 2DG administration or siRNA for PHGDH. AKT stimulates glycolysis and promotes hypertrophy through phosphorylation of P70S6K thereby increasing protein synthesis. Blocking AKT with compound MK-2206 decreases PHGDH abundance and impairs P70S6K phosphorylation which leads to reduced myotube size. Dark red squares indicate where experimental manipulations were applied in this study. Green arrows denote the pathway by which glucose and PHGDH can contribute to muscle size. 2DG, 2-deoxyglucose; MK-2206, AKT blocker.

The first conceptual advance of this study is that glucose is not just a substrate for energy metabolism; it also contributes to cell mass by being a substrate for anabolic reactions both in proliferating C2C12 myoblasts (Hosios et al., 2016) and in post-mitotic, hypertrophying myotubes (Figure 1). Specifically, we found that carbon derived from glucose can be incorporated especially into protein, presumably via a glucose → glycolytic intermediates → amino acid → protein sequence of reactions. This is in line with results reported by Hosios et al., showing that after 14 days, 6% of the total carbon in C2C12 myotubes was derived from glucose (Hosios et al., 2016). Non-essential amino acids are indeed synthesized by human tissues including skeletal muscle (Garber et al., 1976), but it is poorly understood how this is regulated and whether the rate of non-essential amino acid synthesis increases and limits skeletal muscle hypertrophy. The conversion of glucose into biomass may also contribute to the long-term glucose uptake post resistance exercise (Fathinul and Lau, 2009; Marcus et al., 2013) and may help to further explain the beneficial effects of resistance training on metabolic health (Lee et al., 2017).

The second finding of this study is that the inhibition of glycolysis reduces C2C12 and primary myotube size (Figure 2), revealing an association between glycolysis and muscle size. A link between glycolytic flux and growth was first demonstrated by Otto Warburg, who showed that sarcomas, i.e. fast growing tissues, consumed more glucose and synthesized more lactate than normal organs in rats (Warburg et al., 1927). In muscle, the stimulation of muscle hypertrophy in mice through IGF-1 *in vitro* (Semsarian et al., 1999), via gain of *Akt1* (Izumiya et al., 2008), or loss of myostatin (*Mstn*;(Mouisel et al., 2014)) *in vivo* not only results in skeletal muscle hypertrophy, but also increases the glycolytic capacity of the hypertrophying muscles. However, so far there is little data to show whether increased glycolytic flux limits muscle size. Consistent with our findings (Figure 2), the deletion of glycolytic enzymes in flies results in smaller muscle fibers (Tixier et al., 2013). In addition to reducing the supply of glycolytic intermediates as substrates for anabolic reactions, inhibition of glycolysis may also affect the signalling of energy-sensitive signalling molecules such as AMPK (Hardie, 2015) and increase the expression of E3 ubiquitin ligases (Tong et al., 2009). However, we did not find any effect of the glycolysis inhibitor 2DG on activity-related AMPK phosphorylation and E3 ligase expression (Figure 3d,e), suggesting that glycolysis inhibition induced atrophy was likely by attenuation of the rate of protein synthesis. Indeed, phospho-P70S6K levels were increased by IGF-1 and decreased by 2DG. Mechanistically, under low-glucose conditions, the glycolytic enzyme GAPDH prevents Rheb from binding mTORC1, thereby inhibiting mTOR signalling, repressing protein synthesis (Lee et al., 2009).

In relation to the association between glycolytic flux and muscle hypertrophy it is worth noting that glycolytic type 2 fibers typically hypertrophy more after resistance training than less glycolytic type 1 fibers (Andersen and Aagaard, 2000; Kim et al., 2005). This is true even though type 1 fibers have a higher capacity for protein synthesis than type 2 fibers (van Wessel et al., 2010). Future studies should investigate whether the greater hypertrophic potential of type 2 fibers is at least in part, because these fibers can provide more glycolytic intermediates as substrates for anabolic reactions.

The third and fourth findings are that a loss (Figure 4) or gain-of-function (Figure 5) of PHGDH decreases or increases basal and IGF-1-stimulated myotube hypertrophy, respectively. PHGDH was of interest to us because β2-agonist-mediated muscle hypertrophy in pigs increases the expression and protein levels of PHGDH (Brown et al., 2016) and *Phgdh* expression almost doubles when muscle hypertrophy is stimulated through synergist ablation (reanalysis of data from Chaillou et al., 2013)). Our data indicate that PHGDH not only limits cellular proliferation (Possemato et al., 2011) but also the size of post-mitotic cells. This might seem surprising because PHGDH catalyses the de novo synthesis of a non-essential amino acids and so one might assume that the loss of Phgdh would have little effect so long as dietary serine intake is sufficient. However, a complete loss of *Phgdh* is embryonal lethal in mice (Yoshida et al., 2004). *PHGDH* mutations in humans also cause severe inborn diseases, such as Neu-Laxova syndrome (Shaheen et al., 2014), which is associated with atrophic or underdeveloped skeletal muscles (Shved et al., 1985). The importance of PHGDH for normal development and muscle size regulation could be explained through serine’s role as a key metabolite for one-carbon metabolism that is linked to nucleotide and amino acid synthesis, epigenetics and redox defense (Ducker and Rabinowitz, 2017; Reid et al., 2018). A metabolomics analysis showed that PHGDH is required to maintain nucleotide synthesis (Reid et al., 2018), which might affect RNA and ribosome biogenesis in muscle cells. Another mechanism through which PHGDH possibly regulates muscle size is alpha-ketoglutarate. This metabolite is generated downstream of PHGDH and is diminished by 50% upon PHGDH knockdown (Possemato et al., 2011). Mice supplemented with alpha-ketoglutarate are protected against muscle atrophy and increase protein synthesis, inducing muscle hypertrophy (Cai et al., 2016). Future studies should seek to identify the mechanism by which PHGDH contributes to muscle mass in post-mitotic muscle.

We have already mentioned that β2-agonists (Brown et al., 2016) and overload-induced hypertrophy (Chaillou et al., 2013) stimulate the expression of PHGDH, at least temporarily. The fifth finding of this study is that AKT regulates *Phgdh* expression in post-mitotic C2C12 myotubes (Figure 6). In relation to this, future studies are needed to find out whether such increased *Phgdh* expression also occurs in mice where overexpression of AKT causes muscle hypertrophy (Lai et al., 2004) and whether the reduction of PHGDH in these models or in β2-induced muscle hypertrophy reduces muscle size.

Several questions remain unanswered. First, our study only reports *in vitro* data obtained by studying C2C12 and primary myotubes. Whilst both C2C12 myotubes (Peters et al., 2017; Rommel et al., 2001) and mouse muscles respond to IGF-1 with hypertrophy (Musaro et al., 2001), it is unclear whether the *in vivo* hypertrophy involves the same metabolic reprogramming that we report here *in vitro*. Second, whilst 14C is incorporated into protein (Figure 1) it is unclear whether this is because glucose was a substrate for amino acid and protein synthesis or whether proteins became glycosylated (Mariño et al., 2010). Future studies can attempt to use de-glycosylation treatments to verify that carbon from glucose is indeed incorporated into muscle protein.

## 5 Conclusion

In summary, this study provides evidence that glycolysis is important in hypertrophying C2C12 and primary mouse myotubes, reminiscent of cancer-like metabolic reprogramming and that this limits typical myotube size and IGF-1-stimulated muscle hypertrophy.

## Author contributions

H.W., R.J., S.G. conceived the original idea; H.W. R.J., L.S., designed the study; L.S., J.S., B.G., T.H., D.K., I.V., G.W., C.O., performed the experiments; A.M., designed retroviral plasmid; W.H. provided samples; S.V., H.W., R.J. wrote the manuscript; S.V., J.S., B.G., A.M., W.H., H.W., R.T. reviewed and discussed the manuscript.

## Ethics approval

This article does not contain any studies with human participants or animals performed by any of the authors.

## Declaration of Competing Interest

The authors declare no conflict of interest

## Acknowledgements

B.M.G. was supported by fellowships from the Novo Nordisk Foundation (NNF19OC0055072) & the Wenner-Gren Foundation, an Albert Renold Travel Fellowship from the European Foundation for the Study of Diabetes and an Eric Reid Fund for Methodology from the Biochemical Society. A.D.M. was funded initially by Sarcoma UK (grant number SUK09.2015), then supported by funding from Postdoctoral Fellowship Program (Helmholtz Zentrum München, Germany) and currently by Cancer Research UK.

## Notes

### Competing Interest Statement

The authors have declared no competing interest.

